# Six months of physical inactivity is insufficient to cause chronic kidney disease in C57BL/6J mice

**DOI:** 10.1101/2024.08.29.610415

**Authors:** Precious C. Opurum, Stephen T. Decker, Deborah Stuart, Alek D. Peterlin, Venisia L. Paula, Piyarat Siripoksup, Micah J. Drummond, Alejandro Sanchez, Nirupama Ramkumar, Katsuhiko Funai

**Affiliations:** Diabetes & Metabolism Research Center, University of Utah, Salt Lake City, Utah, USA; Department of Nutrition and Integrative Physiology, University of Utah, Salt Lake City, Utah, USA; Molecular Medicine Program, University of Utah, Salt Lake City, Utah, USA; Division of Nephrology & Hypertension, School of Medicine, University of Utah, Salt Lake City, Utah, USA; Department of Physical Therapy & Athletic Training, University of Utah, Salt Lake City, Utah, USA; Division of Urology, Department of Surgery, University of Utah School of Medicine, Salt Lake City, Utah, USA; Huntsman Cancer Institute, Cancer Hospital, Salt Lake City, Utah, USA

## Abstract

Chronic kidney disease (CKD) is a progressive disorder marked by a decline in kidney function. Obesity and sedentary behavior contribute to the development of CKD, though mechanisms by which this occurs are poorly understood. This knowledge gap is worsened by the lack of a reliable murine CKD model that does not rely on injury, toxin, or gene deletion to induce a reduction in kidney function. High-fat diet (HFD) feeding alone is insufficient to cause reduced kidney function until later in life. Here, we employed a small mouse cage (SMC), a recently developed mouse model of sedentariness, to study its effect on kidney function. Wildtype C57BL/6J male mice were housed in sham or SMC housing for six months with HFD in room (22°C) or thermoneutral (30°C) conditions. Despite hyperinsulinemia induced by the SMC+HFD intervention, kidneys from these mice displayed normal glomerular filtration rate (GFR). However, the kidneys showed early signs of kidney injury, including increases in Col1a1 and NGAL transcripts, as well as fibrosis by histology, primarily in the inner medullary/papilla region. High-resolution respirometry and fluorometry experiments showed no statistically significant changes in the capacities for respiration, ATP synthesis, or electron leak. These data confirm the technical challenge in modeling human CKD. They further support the notion that obesity and a sedentary lifestyle make the kidneys more vulnerable, but additional insults are likely required for the pathogenesis of CKD.

## INTRODUCTION

Obesity and its comorbidities, such as diabetes and hypertension, are key risk factors for chronic kidney disease (CKD) (1, 2). The global surge of individuals with end-stage renal disease (ESRD) who require kidney transplants or dialysis is reaching an epidemic scale. The number of people with the early stages of CKD (stages 1 to 4) is projected to be 50 times greater than those with ESRD (stage 5) (3, 4). The Centers for Disease Control and Prevention (CDC) estimates that 37 million adults in the US, or 15% of the population, have CKD, with an average life expectancy of 8 years after diagnosis for patients on dialysis. Forty percent of the population with significantly impaired kidney function (not on dialysis) are unaware of their condition (5). Currently, there are no therapeutics to directly enhance kidney function in CKD patients (3). Rather, most medications used clinically prevent the progression of CKD (such as SGLT2 inhibitors)(6, 7). A recent study shows that semaglutide lowers the incidence of CKD, though this is likely mostly secondary to reduced adiposity (8, 9).

Highly heterogenous systemic and metabolic factors contribute to the pathogenesis of CKD. Obesity doubles the chance of developing risk factors for CKD and directly impacts the progression of CKD and ESRD (10). Obesity may cause renal dysfunction and damage even without diabetes or hypertension via adipose and hemodynamic-mediated effects on kidneys (11). Inadequate physical activity independently contributes to the development of CKD beyond its impact on obesity (12). Nevertheless, the precise mechanisms by which obesity or sedentary behavior promotes CKD remain unknown (13).

Currently, mechanistic studies for CKD are highly limited by the lack of appropriate murine models. Widely used mouse models of reduced kidney function, such as ischemic reperfusion and unilateral ureteral obstruction, are more accurately described as models for acute kidney injury (AKI). In contrast, chronic models, such as adenine diet feeding, induce systemic and renal stress that is different from conditions seen in human CKD (14). While many advances in our understanding have been made with these models, it is difficult to imagine that they fully recapitulate the underlying processes that occur in humans (15). Nonetheless, it is important to point out that AKI may be an integral component of the pathogenesis of CKD. Several studies show that AKI accelerates the course of CKD and that CKD increases the susceptibility of kidneys to subsequent AKI, suggesting a bidirectional relationship between AKI and CKD (16, 17).

Our team recently developed a new model for physical inactivity in mice by employing a cage size reduction model used in rats (18). Unlike many traditional models of physical inactivity, such as hindlimb unloading and denervation, which are acute challenges that trigger unnatural endocrine responses in mice, the small mouse cage (SMC) promotes metabolic inflexibility without increasing circulating cortisol levels (19). For the current study, we employed SMC in conjunction with HFD feeding for six months to test our hypothesis that physical inactivity makes mice with diet-induced obesity more prone to develop CKD (20). We performed these studies both at room temperature and at thermoneutrality. The goal of these studies was to establish a murine model of CKD that does not rely on injury, toxin, or gene deletion to reduce kidney function. To make the model widely useful to the kidney research community, we chose to study wild-type mice on the C57BL/6J background. We also performed in-depth phenotyping of kidney functional and metabolic parameters to evaluate the independent contributions of obesity and physical inactivity to the development and progression of CKD.

## MATERIALS AND METHODS

### Study Approval

Experiments on animals were performed per the Guide for the Care and Use of Laboratory Animals of the National Institutes of Health. All animals were handled according to approved University of Utah Animal Use and Care Committee (IACUC) protocols (#1633).

### Animals

Male C57BL/6J mice from Jackson Laboratory (Strain# 000664) were maintained with a standard chow diet or 42% high-fat diet (HFD; Envigo TD.88137) for 24 weeks at room temperature (22°C) or thermoneutrality (30°C). Mice started their intervention at 8 weeks of age for experiments at room temperature and 20 weeks for experiments at thermoneutrality. Mice were placed in standard housing, and all animals had access to food, water, and ad libitum throughout the study duration and were fasted 3 hours before tissue collection. Body weights, food, and water intake were monitored daily with weekly measurements. Body composition was determined using the Bruker Minispec NMR (Bruker, Billerica, MA) at 8-week intervals (0, 8,16, and 24 weeks). After 24 weeks of chow or HFD, mice were fasted for 3 hours before being euthanized using a lethal mix of ketamine and xylazine.

### Small Mouse Cage (SMC)

Our SMC is designed as a rectangular box produced from acrylic plastic and made at the University of Utah’s Machine Shop Core (Fig 1A). The current SMC design was modified and further developed from our previous SMC design used by Siripoksup *et al* (19). The bedding is laid up to three-quarters of the height of the cage, limiting space for mice to move around. The SMC mice were placed inside a standard housing cage and were paired with a sham mouse to prevent social isolation-induced stress; a clear acrylic divider separated each mouse to prevent the sham mice from consuming food and water from the SMC cage (Fig 1B).

**Figure 1.**
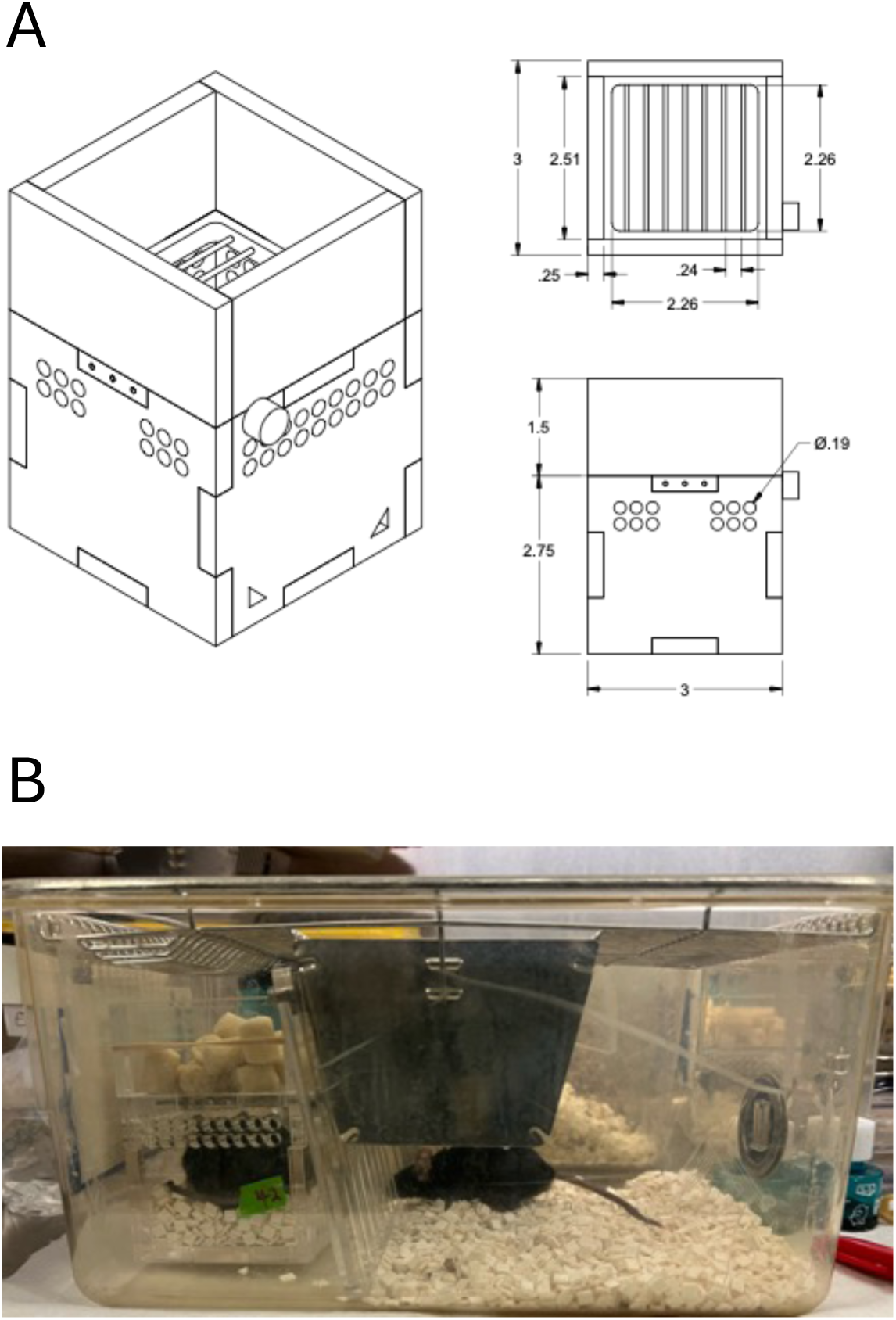
Design and setup of the Small Mouse Cages (SMC). (A) Dimensions of the cages used in this study; (B) Representative image of the paired housing setup. One mouse is placed in the SMC, with *ad libitum* access to food (top) and water (not visible). The other paired mouse is placed in the open side of the cage, separated by an acrylic barrier.

### Glomerular filtration rate (GFR) Measurements

GFR was measured using the renal clearance of fluorescein isothiocyanate-labeled sinistrin (FITC-sinistrin) from Medibeacon (Medibeacon GMBH, Mannheim, Germany), as previously described (21). During isoflurane anesthesia, a patch of dorsal skin was shaved, and a non-invasive clearance (NIC-Kidney device: Medibeacon) transcutaneous monitor was attached and monitored background fluorescence for 10 minutes. FITC-sinistrin was administered retro-orbitally (7.5 mg/100 g body weight). GFR was measured using FITC-sinistrin clearance and fluorescence monitoring for the next hour. The kinetics of fluorescence degradation were utilized to determine GFR using the manufacturer’s software, following their standard guidelines.

### Serum Insulin and Blood urea nitrogen (BUN)

Blood was drawn from the submandibular (facial) vein before anesthesia, placed in tubes containing EDTA, and kept on ice. Blood was centrifuged at 2400 g for 10 minutes at 4 °C. The supernatant (serum) was collected, transferred to a fresh tube, and stored at −80°C until analysis. Serum insulin levels were measured with a Mouse Insulin ELISA kit (Cat# 90080 Crystal Chem, Chicago, Illinois). Blood urea nitrogen (BUN) was measured using a commercially available quantitative colorimetric analysis according to the manufacturer’s protocol (Quantichrom, Urea Assay Kit Cat# DIUR-100, BioAssay Systems, Hayward, CA).

### Urine collection

Mice were placed in an MMC100 Metabolic Cage, and urine was collected every 24 hours for 48 hours at the end of the study. Body weight, food, and water intake were all observed throughout the experiment. Mice were fed with standard chow or an HFD diet and converted into a gel diet with free access to water. Urine samples were centrifuged at 21,000 g for 15 minutes, then aliquoted and stored at −80°C until the assay. Urine was measured using QuantiChrom™ Creatinine Assay Kit, BioAssay Systems, Hayward, CA (Cat# DIUR-500), and albuminuria was quantified using Albuwell M (Mouse Albumin ELISA, Ethos Biosciences, Philadelphia, PA) according to manufacturer instructions. Urine ACR was calculated as the ratio of albumin to creatinine.

### Histology

Kidney samples were fixed in 10% formaldehyde for 24-48 hours and transferred to the ARUP Research Histology Core Laboratory at the University of Utah. Kidney slices were fixed in paraffin, sectioned at 4 μm, and stained with Masson’s trichrome to detect morphological changes and fibrosis. Trichrome-stained Kidney sections were imaged using Zeiss AxioScan.Z1 (ZEISS, Oberkochen, Germany) slide scanner at the University of Utah Imaging core.

### Real-time polymerase chain reaction (RT-qPCR)

For real-time quantitative polymerase chain reaction (qPCR) investigations, whole mouse kidney tissues were lysed in 1 ml of TRIzol (Thermo Fisher Scientific, USA), and RNA was extracted using conventional methods. We used the iScript cDNA Synthesis Kit (Bio-Rad, Hercules, CA) to reverse transcribe total RNA. To assess the expression of kidney damage biomarkers: Kidney Injury Molecule 1 (KIM-1), Collagen I (Col1a1), Collagen IV (Col4a1), and Neutrophil gelatinase-associated lipocalin (NGAL), pre-validated primer sequences were taken from either the primer bank or previously published papers. All mRNA levels were standardized to RPL32 with SYBR Green reagents (Thermo Fisher Scientific, USA), and 384 well plates were loaded and sent to the University of Utah Genomics Core labs for qPCR.

### Kidney mitochondrial isolation

Kidney tissues were minced in ice-cold mitochondrial isolation medium (MIM) buffer [300 mM sucrose, 10 mM HEPES, 1 mM EGTA, and 1 mg/ml bovine serum albumin (BSA) (pH 7.4)] and carefully homogenized with a Teflon-glass pestle. The homogenate was centrifuged at 800 g for 10 minutes at 4 °C. The supernatant was transferred to another tube and centrifuged at 1,300 g for 10 minutes. The supernatant was transferred to another tube and centrifuged at 10,000 g for 10 minutes at 4 °C. The pellet was resuspended in an MIM buffer, depending on its size. Protein concentrations were assessed with Pierce™ BCA Protein Assay Kits (# 23225, Thermo Fisher Scientific, USA).

### High-resolution respirometry and fluorometry

Mitochondrial O2 consumption was evaluated using the Oroboros Oxygraph O2K (Oroboros Instruments, Innsbruck, Austria) as previously described (22). The isolated mitochondria were introduced to the oxygraphy chambers containing 2 ml of Buffer Z (105 mM MES potassium salt, 30 mM potassium chloride (KCl), 10mM monopotassium phosphate (KH_2_PO_4_), 5mM magnesium chloride (MgCl_2_), and 0.5 mg/ml BSA). Respiration was assessed by adding the following substrates: 0.5 mM malate, 5 mM pyruvate, 5 mM glutamate, 2 mM ADP, 10 mM succinate, and 1.5 μM FCCP (Carbonyl cyanide-p-trifluoromethoxyphenylhydrazone).

ATP production was fluorometrically quantified with a Horiba Fluoromax 4 (Horiba Scientific, USA) by enzymatically linking it to NADPH synthesis, as previously described (22). ATP generation was assessed with 0.5 mM malate, 5 mM pyruvate, 5 mM glutamate, 10 mM succinate, and successive 2, 20, and 200 μM ADP additions.

All hydrogen peroxide (*J*H_2_O_2_) studies were conducted in buffer Z supplemented with 10 mM Amplex Ultra Red (Invitrogen, Waltham, MA) and 20 U/ml CuZn SOD (superoxide dismutase) as previously described (22). In brief, isolated mitochondria were added to 1 ml of assay buffer containing Amplex Ultra Red, which creates a fluorescent product when oxidized by H_2_O_2_. H_2_O_2_ emission was measured in the presence of 10 mM succinate. The fluorescent product’s appearance was measured using a Horiba Fluoromax 4 fluorometer (excitation/emission 565/600). Auranofin and BCNU (bis-chloroethylnitrosourea) were used to inhibit thioredoxin and glutathione reductase, respectively, which are known to affect H_2_O_2_ measurements.

### Western blot analyses

Whole kidney tissue samples were homogenized in cold RIPA lysis buffer (#89901, Thermo Fisher Scientific, USA) containing protease inhibitor (#78446, Thermo Fisher Scientific, USA), incubated at 4°C for 10 min, and centrifuged at 12,000 *g* for 15 min at 4°C, and the supernatant was collected into a fresh tube. The BCA Protein Assay Kit (#23225, Thermo Fisher Scientific, USA) was used to measure the protein concentration in the supernatant. Western blotting was carried out as previously described (23) with slight modifications. In summary, for electrophoresis, appropriate volumes of protein were combined with 4x Laemmli sample buffer supplemented with 10% β-mercaptoethanol (#1610710, Bio-Rad, Hercules, CA) and loaded onto a Bio-Rad 4%–20% gradient SDS-polyacrylamide gel. After that, the proteins were transferred from the gel onto membranes made of polyvinylidene fluoride (Thermo Fisher Scientific, USA). The membranes were blocked for one hour at room temperature using 5% BSA in Tris-buffered saline containing 0.1% Tween 20 (TBS-T), and then the kidney samples were incubated overnight with specific primary antibodies to probe for protein abundance of OXPHOS protein complexes (STN-19467, Abcam), 4-HNE (ab48506, Abcam), Actin (A2066, Sigma-Aldrich), Citrate synthase (AB9660, Abcam). Following multiple TBS-T washes, the membranes were incubated for 1 hour at room temperature with appropriate secondary antibodies diluted 1:5,000 in 5% BSA. After multiple washes in TBS-T and one final rinse in TBS. Membranes were exposed with Western Lightning Plus-ECL (PerkinElmer) and imaged using a ChemiDoc Imaging System (Bio-Rad, Hercules, CA). The images were quantified using Image Lab software (Bio-Rad)

### Statistical analyses

All results are expressed as means ± SEM. For statistical comparisons, one- or two-way ANOVA followed by Tukey’s multiple comparison tests were performed using GraphPad Prism v10.3.0 software (GraphPad Software, Boston, MA). The criterion for significance cutoff was P ≤ 0.05.

## RESULTS

### Physical inactivity does not reduce kidney function in mice housed at room temperature

Our pilot studies showed that 8 and 16-week SMC+HFD interventions were insufficient to reduce GFR (data not shown). We then examined a 24-week intervention with the goal of designing a murine CKD model that is relatively simple, and sufficiently short in duration to minimize the impact of aging on kidney function. For experiments at room temperature, 8-week-old mice were fed standard chow (Sham Chow), HFD (Sham HFD), or HFD in SMC (SMC HFD). Mice on a chow diet only marginally gained weight during the 24-week period, while mice in both Sham HFD and SMC HFD groups gained weight (∼40 g) during the 24-week intervention (Fig 2A). The body masses between Sham HFD and SMC HFD was not the same throughout the 24-week period, as the Sham HFD group gained weight more rapidly and reached ∼40 g faster than the SMC HFD group. There were no differences in lean mass among the groups (Fig 2B), as variations in fat mass fully accounted for the differences in body mass (Fig 2C). We previously showed that SMC promotes hyperinsulinemia (19). In the current study, HFD feeding also led to hyperinsulinemia, and the SMC intervention had an additive effect of increasing circulating insulin further (Fig 2D). These data suggest that HFD feeding combined with sedentary behavior facilitated by SMC housing had additive effects on systemic metabolic stress.

**Figure 2.**
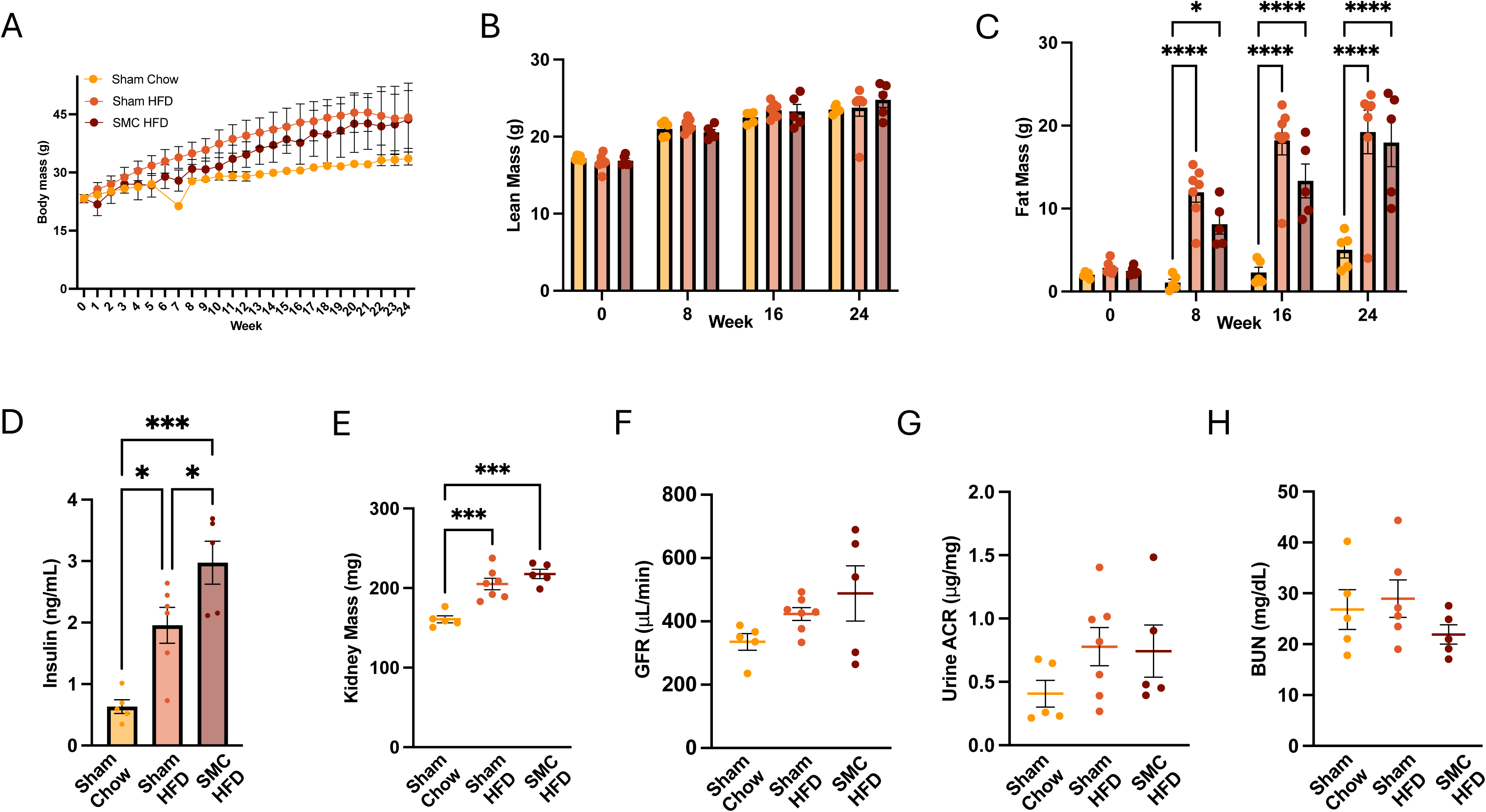
Combined SMC and HFD intervention in mice housed at room temperature. (A) Whole body, (B) Lean, and (C) Fat mass of the chow-fed (sham chow), high-fat diet-fed (Sham HFD), and small mouse caged (SMC) and high-fat diet-fed animals (SMC HFD), (D) Fasting plasma insulin levels and (E) Kidney mass. Kidney function analysis via (F) Glomerular filtration rate, (G) Urine albumin to creatinine ratio, and (H) Blood urea nitrogen at room temperature (22°C), n=5-10. P value *<0.05, *** <0.005. **** <0.0005. HFD ⎯ High-fat Diet; SMC ⎯ Small mouse cage; RT ⎯ Room temperature; GFR – Glomerular Filtration Rate.

To evaluate the effects of HFD feeding or HFD feeding combined with physical inactivity on kidney function, we next examined the kidneys of these mice. HFD feeding, but not SMC, increased kidney mass during the intervention (Fig 2E). To evaluate the effects of HFD and SMC interventions on kidney function, we assessed glomerular filtration rate (Fig 2F), albuminuria (Fig 2G), and blood urea nitrogen (Fig 2H). However, none of these parameters were statistically different among the three groups.

### Physical inactivity intervention combined with HFD feeding does not induce kidney dysfunction in mice housed at thermoneutrality

To follow up with the experiments in Fig 2, we performed an additional SMC/HFD intervention study with two modifications. First, mice were housed at thermoneutrality rather than at room temperature. Mice housed at room temperature experience non-shivering thermogenesis to maintain core body temperature, thus increasing energy expenditure. It is conceivable that such an increase in energy expenditure may counteract the adverse metabolic effects of SMC housing and HFD feeding. Second, we began the 24-week intervention in 20-week-old mice.

Our rationale for choosing older mice was based on the notion that mice are still growing at 8 weeks, which also requires energy and could attenuate metabolic disturbances that would otherwise be observed in fully-grown adults. We decided against extending the SMC/HFD interventions further because by the end of the 24 weeks, the mice would be close to 1 year old, and aging effects might start influencing kidney function. Additionally, we aimed to create a murine CKD model that was relatively easy to recapitulate, and we felt that extending the intervention to 9-12 months was counterproductive to this goal.

To determine the effects of HFD feeding and physical inactivity with or without thermoneutrality, mice were divided into four groups: mice fed standard chow at room temperature (Sham Chow RT), mice fed standard chow at thermoneutrality (Sham Chow TN), mice fed HFD at thermoneutrality (Sham HFD TN), and mice fed HFD and housed in SMC at thermoneutrality (SMC HFD TN). Body weights for Sham Chow RT and Sham Chow TN groups did not change during the 24-week intervention (Fig 3A). Similar to our findings from Fig 2, body weights in Sham HFD TN and SMC HFD TN groups were similar at the end of the 24-wk intervention, though SMC HFD TN groups gained weight more slowly than the Sham HFD TN group (Fig 3A). Lean masses were not different among the four groups at the end of the intervention (Fig 3B), and differences in body weights were again explained by differences in fat mass (Fig 3C). HFD and SMC+HFD interventions induced hyperinsulinemia (Fig 3D), though there were no differences in fasting circulating insulin between Sham HFD TN and SMC HFD TN groups.

**Figure 3.**
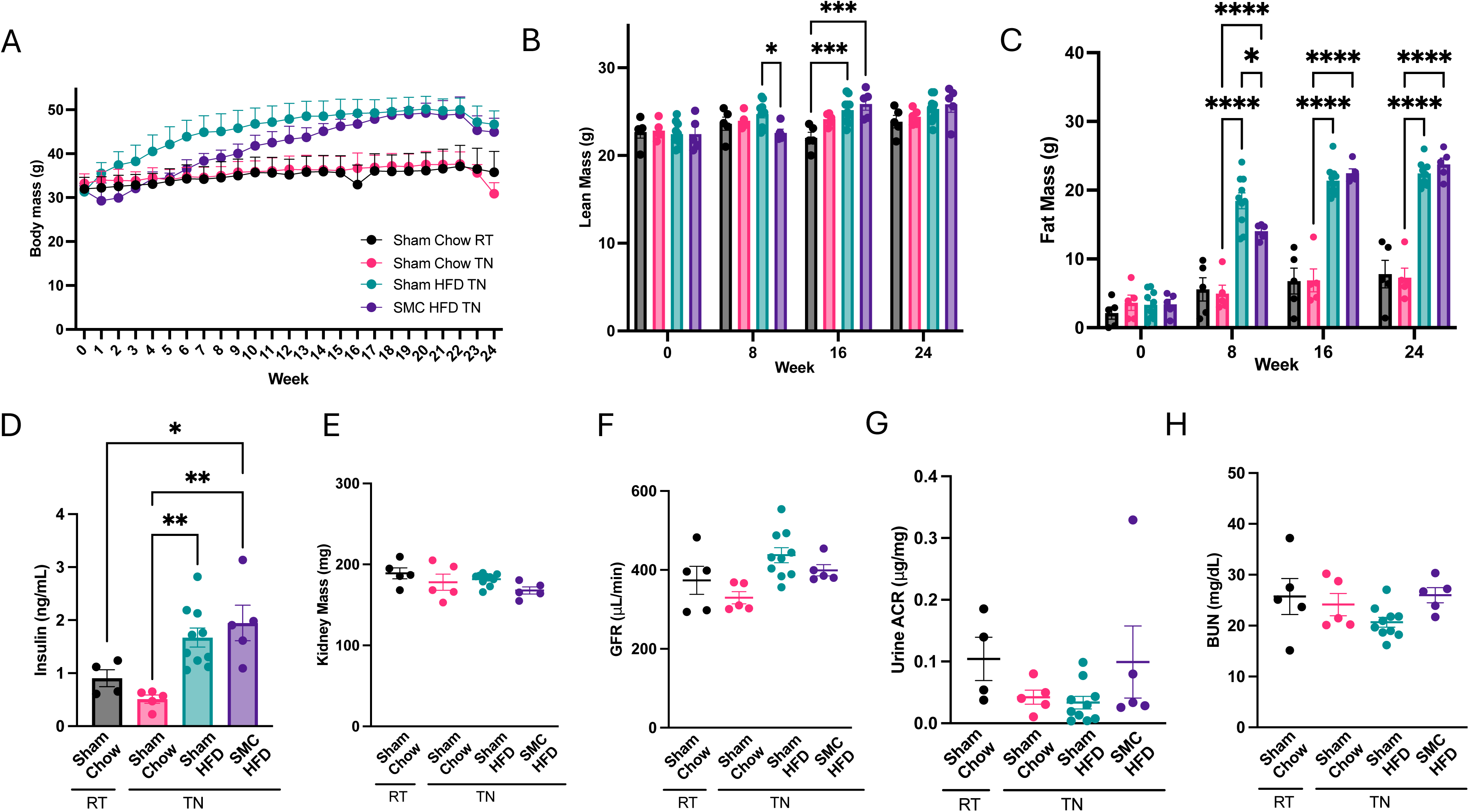
Combined SMC and HFD intervention in mice housed at thermoneutrality. (A) Whole body (B) Lean, and (C) Fat mass of the chow-fed at RT (sham Chow RT) and Thermoneutrality (Sham Chow TN), high-fat diet-fed at TN (Sham HFD TN), and small mouse caged (SMC), high-fat diet-fed at TN animals (SMC HFD TN). (D) Fasting plasma insulin levels and (E) Kidney mass. Kidney function analysis via (F) Glomerular filtration rate, (G) Albuminuria, (H) Blood urea nitrogen at thermoneutral conditions (30°C), n=5-10. P value *<0.05, *** <0.005. **** <0.0005. HFD ⎯ High-fat Diet; SMC ⎯ Small mouse cage; RT ⎯ Room temperature; TN ⎯ Thermoneutrality; GFR – Glomerular Filtration Rate.

To determine the effects of these interventions on kidney function, we then examined the kidneys of these mice. There were no differences in kidney mass (Fig 3E). We also evaluated kidney functions, but GFR (Fig 3F), albuminuria (Fig 3G), and blood urea nitrogen (Fig 3H) were not influenced by the interventions. Thus, 6-month HFD and HFD+SMC interventions in room temperature or thermoneutrality are insufficient to cause CKD in C57BL/6J mice.

### Physical inactivity promotes fibrotic programming in kidney

While GFR is the hallmark of kidney function and its reduction is a defining feature of CKD, the progression to CKD is often associated with earlier changes in various aspects of the kidney that precede a decline in GFR. One of these changes is the onset of fibrotic programming (24). Indeed, mRNA levels of Col1a1 and NGAL were elevated in the SMC HFD TN group compared to all other groups (Fig 4A). Histological analyses of these kidneys revealed increased fibrosis, particularly in the kidney’s inner medullary/papilla (Fig 4B-E) and less so in the cortex region (Fig 4F-I). Trichrome blue staining in the cortex region was only marginally greater in the SMC HFD TN group compared to others, and therefore inconclusive as to whether the SMC/HFD intervention promoted fibrosis in the cortex. Nevertheless, these observations may indicate changes that could make the kidneys more susceptible to reduced function if an additional insult occurs. No signs of fibrosis were observed in the kidneys of mice housed at room temperature (Supplemental Fig S1).

**Figure 4.**
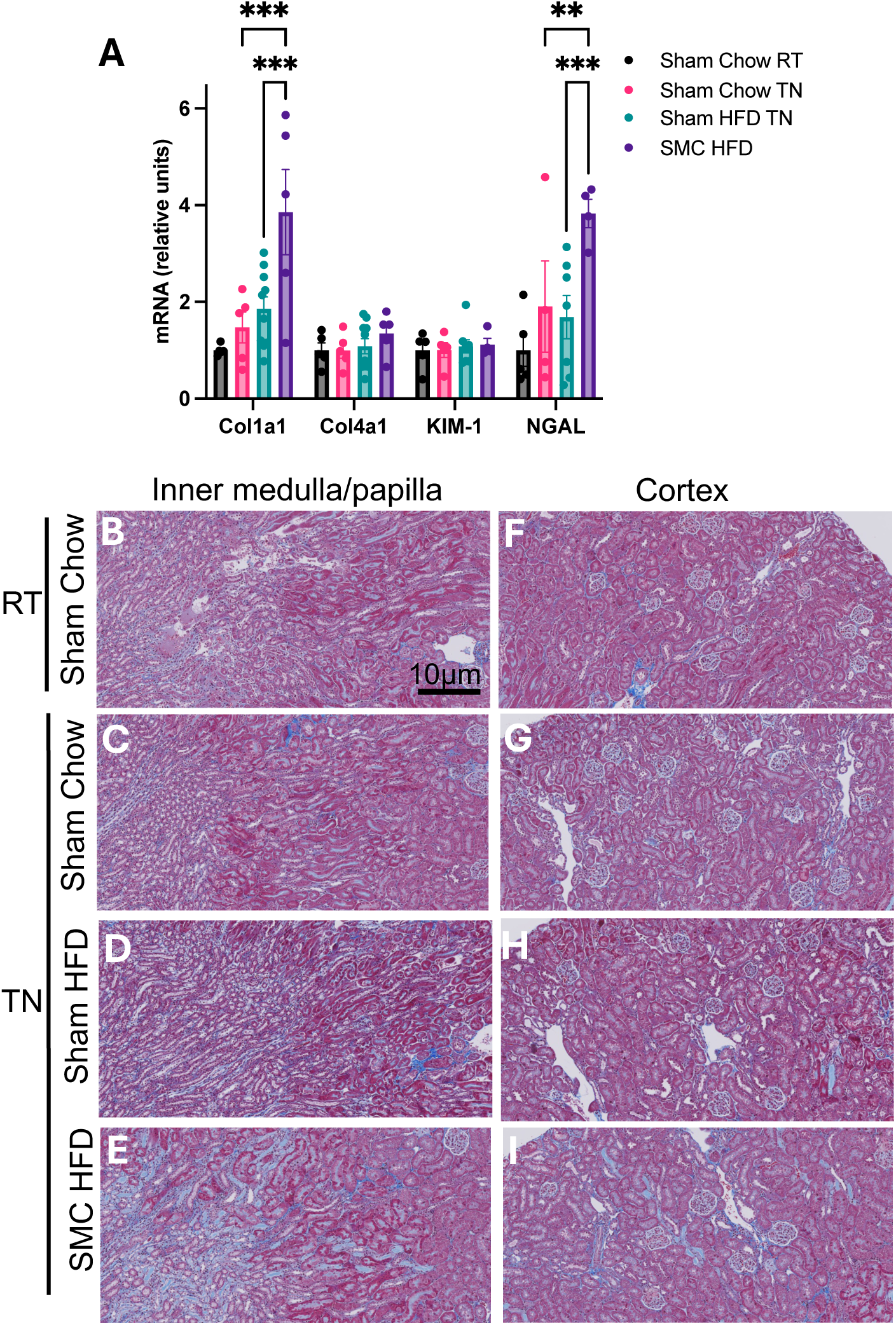
Combined SMC and HFD intervention induce mild kidney fibrosis. (A) Kidney damage markers: collagen I & IV, kidney injury marker-1 (KIM-1), and neutrophil gelatinase-associated lipocalin (NGAL) relative expression. Trichrome-stained kidney inner medullary/papilla (B-E) and cortex (F-I) sections from sham and SMC mice fed chow or HFD housed at room temperature or thermoneutral conditions. P value =0.05 (P value *<0.05, ** <0.005, *** <0.0005). Images are representative of n=5-10 per group. Magnification at x20 zoomed at 30% using Zeiss AxioScan.Z1 slide scanner. HFD ⎯ High-fat Diet; SMC ⎯ Small mouse cage; RT ⎯ Room temperature; TN ⎯ Thermoneutrality.

### Physical inactivity does not negatively influence mitochondrial bioenergetics in kidney

Kidney function largely depends on the proximal tubule epithelial cells critical for nutrient reuptake. These cells contain an extraordinarily high mitochondria density, only second to retinal ganglion cells (25, 26). Defects in mitochondrial oxidative phosphorylation have been implicated in the pathogenesis of CKD, particularly with respect to reduced capacity for ATP production, as well as increased propensity for electron leak and oxidative stress (27). Because changes in mitochondrial energetics likely preceded reduced GFR, we assessed mitochondrial bioenergetic properties in the kidneys of our mice.

Any alterations in mitochondrial energetics could be due to changes in mitochondrial density in the cells. We assessed the abundance of representative subunits from respiratory complexes (Complex I, II, III, IV, and V) as well as a matrix protein citrate synthase (Fig 5 A-C). SMC and/or HFD intervention had no effect on the abundance of these enzymes, suggesting that mitochondrial density was not altered with these interventions. We then assessed kidney mitochondrial oxidative capacity by high-resolution Oroboros respirometry (Fig 5D). No differences were observed in mitochondrial respiration. Using high-resolution fluorometry, we also quantified the capacity for mitochondrial ATP production under increasing energetic demand assessed with increasing concentrations of ADP (Fig 6A). There were trends for thermoneutrality to reduce ATP production capacity, but these differences were not statistically significant. Importantly, HFD or SMC/HFD intervention had no influence on ATP production rates. We also quantified oxygen consumption under variable ADP concentrations, which also showed no differences among the groups (Supplemental Fig S2A).

**Figure 5.**
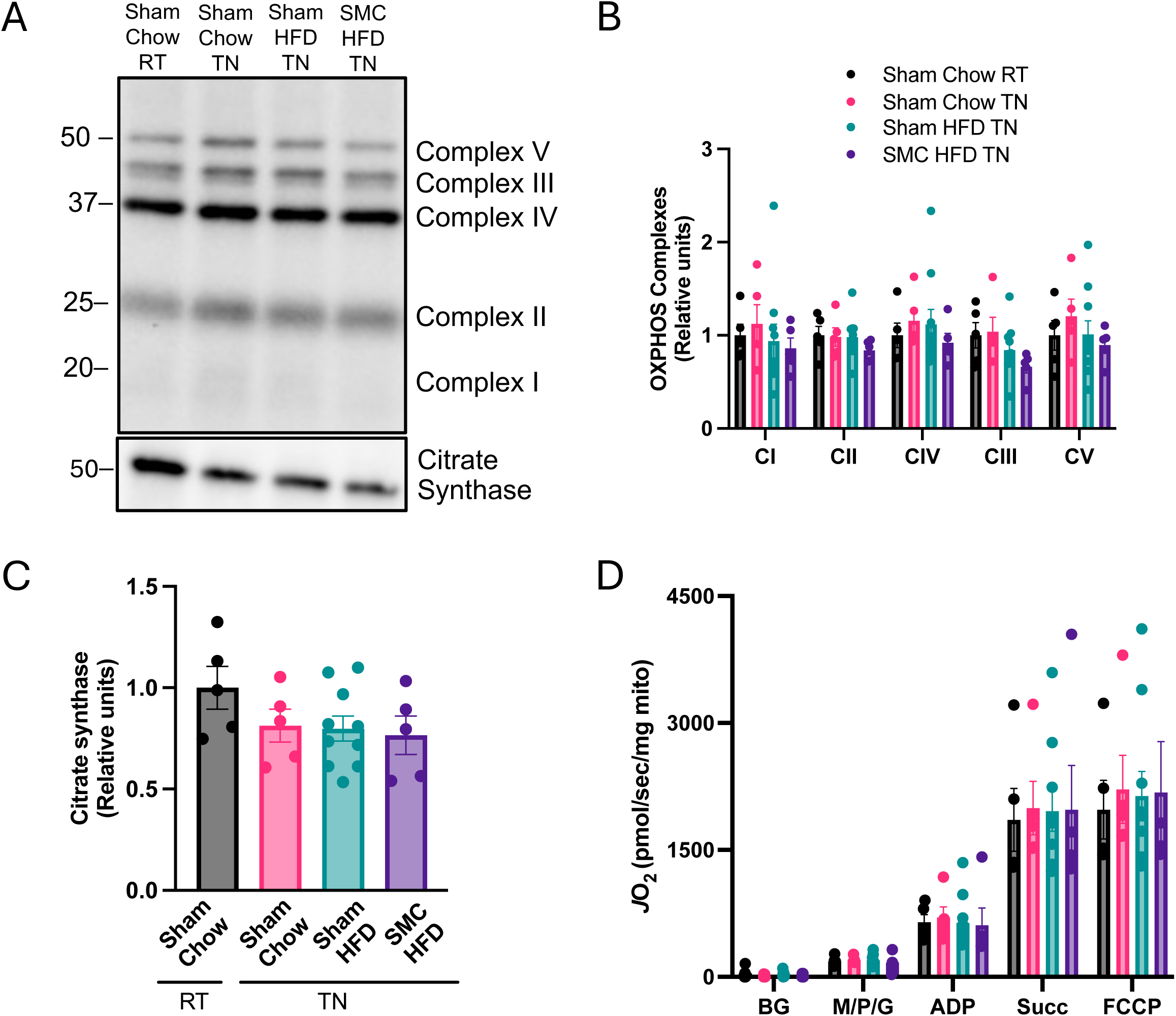
Mitochondrial bioenergetic phenotyping in kidney. (A) Western blots and (B) quantification for OXPHOS complexes I - V and (C) citrate synthase in whole kidney lysate. (D) O_2_ utilization using enriched kidney mitochondria fraction with glycolytic substrates (2 mM ADP, 0.5 mM malate, 5 mM pyruvate, 10 mM succinate, 1 μM carbonyl cyanide-*p*-trifluoromethoxyphenylhydrazone (FCCP) n=5-10 per group. HFD ⎯ High-fat Diet; SMC ⎯ Small mouse cage; RT ⎯ Room temperature; TN ⎯ Thermoneutrality.

**Figure 6.**
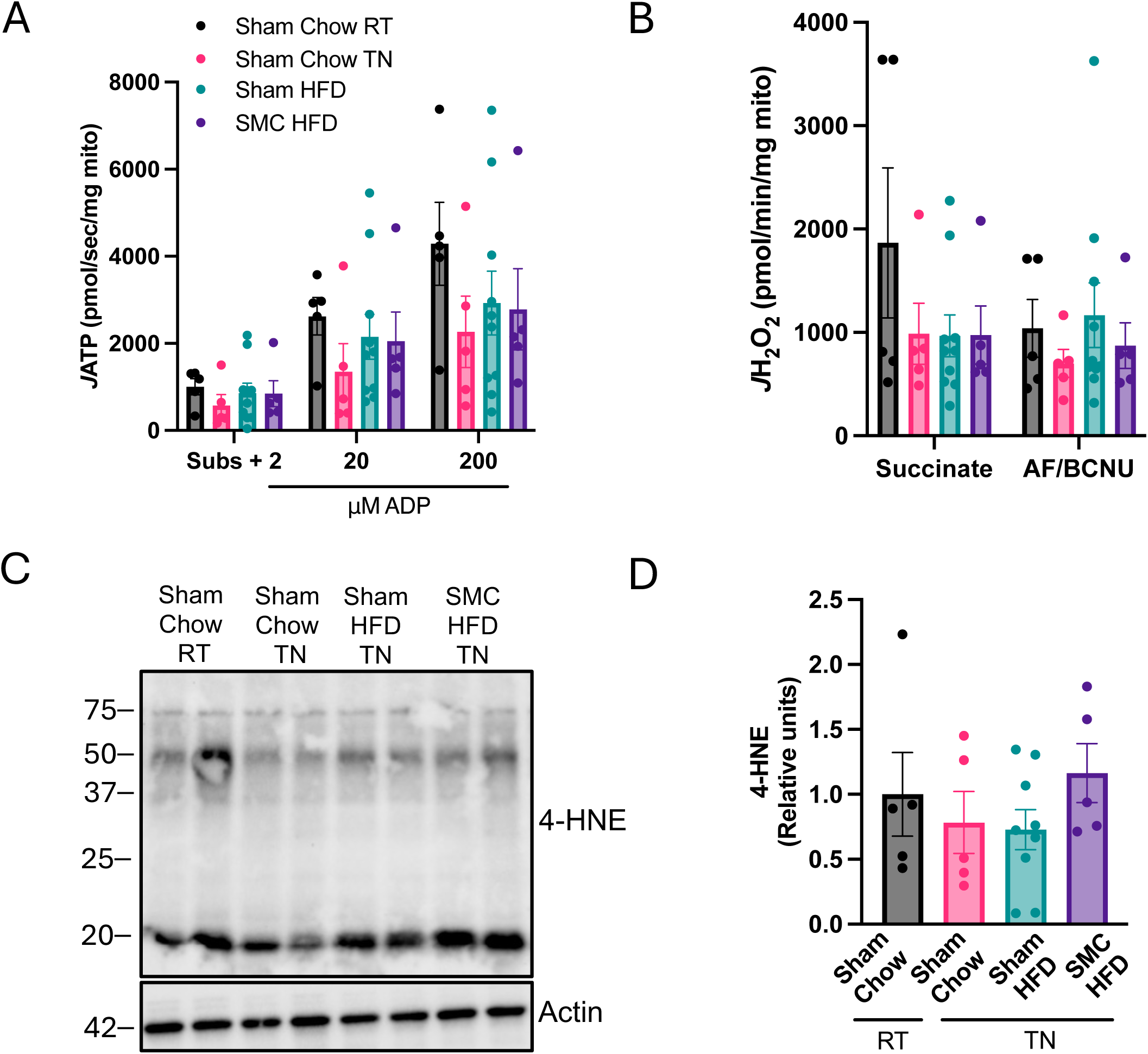
Mitochondrial ATP production and electron leak. (A) ATP production with ADP titrations, and (B) H_2_O_2_ emission using enriched kidney mitochondria fraction. (C) Western blots and (D) quantification for 4-hydroxynonenal (4-HNE) and actin in whole kidney lysate of sham and SMC mice at room temperature and thermoneutrality, n=5-10 per group. HFD ⎯ High-fat Diet; SMC ⎯ Small mouse cage; RT ⎯ Room temperature; TN⎯ Thermoneutrality; BG-Background; ADP-Adenosine 5’-diphosphate; AF-Auranofin; BCNU-Bis-chloroethylnitrosourea/carmustine.

Electrons escaping from the electron transport chain can react with molecular oxygen and promote oxidative stress. Because a very small fraction of electrons normally escape from OXPHOS, increases in electron leak can happen in the absence of a measurable decrease in oxygen consumption or ATP production. Using high-resolution fluorometry, we quantified mitochondrial electron leak in kidneys from our mice (Fig 6B). However, there were no differences in the electron leak between the groups. We also confirmed that these interventions had no influence on oxidative stress by quantification of lipid carbonyl 4-HNE (Fig 6C and D). Findings were similar for kidneys from mice housed at room temperature (Supplemental Fig S2B-F, S3A-C). Thus, our combined SMC and HFD interventions in the current study are insufficient to robustly alter kidney mitochondrial bioenergetics in C57BL/6J mice.

## DISCUSSION

In this study, we sought to establish a murine model of physical activity that can be used to study the pathogenesis of CKD. However, six months of SMC intervention combined with HFD feeding was insufficient to induce CKD in mature and middle-aged C57B6/J mice. Nonetheless, SMC intervention induced changes that indicated an onset of fibrotic programming in the kidney, suggesting that the SMC intervention provides additional metabolic insult not observed with HFD feeding alone. To that effect, SMC+HFD intervention may be employed with other genetic or environmental insults to study CKD. Our data, which shows that six months of physical activity and HFD feeding were insufficient to reduce GFR, is also consistent with the idea that additional insults, such as elevated blood pressure, diabetes or AKI, may play an important role in accelerating the pathogenesis of CKD.

Renal proximal tubular inflammatory damage from HFD feeding may contribute to chronic kidney disease (28, 29). However, HFD feeding alone may not affect renal function in the short-term (30). Physical inactivity is a major risk factors for human CKD, but very little is known regarding its mechanism (31). Current clinical practice guidelines from KDIGO recommend moderate-intensity physical activity for a cumulative duration of 150 minutes per week to reduce the progression of CKD (32). The kidney is one of the major organs that can be directly modulated by physical activity (33), and the notion that CKD patients benefit from greater physical activity is well supported (32, 34). These benefits include better eGFR and prevention of muscle wasting, a major complication associated with mortality in CKD patients. Physical activity also reduces the risk of developing obesity, diabetes, and hypertension, all of which are important risk factors for CKD (35). However, several studies argue that the type and intensity of the exercise beneficial for CKD patients may vary depending on the rigor of the activity in question and disease state (36, 37). The observed changes in kidney histological structures, including an increase in inner medullary/papilla fibrosis supported by high Col1a1 and NGAL in the physically inactive HFD feed group at thermoneutrality, are consistent with the idea that an additional insult is necessary to precipitate the effects of HFD on kidney proximal tubules. Increased fibrotic biomarkers Col1a1 & NGAL is predictive of early signs of CKD (38, 39) and may be related to a failed attempt at injury repair and regeneration processes, indicating the kidney’s adaptation to preserve normal function (40, 41). Fibrotic programming is strongly correlated with CKD progression, which typically precedes an apparent reduction in GFR (24). Decleves *et al*. (42) showed that HFD-associated inflammatory and fibrotic markers appear before albuminuria in mice fed 60% HFD for 12 weeks. This suggests that we expect even slower progression to proteinuria with no variations in BUN levels (43), which aligns with our results.

In addition to the SMC intervention and HFD feeding, we also introduced two additional variables to examine whether these changes could make mice more susceptible to CKD. First, we housed mice at thermoneutrality. Mice activate non-shivering thermogenesis at room temperature, increasing energy expenditure. Housing mice at thermoneutrality lowers metabolic rates and makes them more prone to developing metabolic diseases (44–47). Second, we also began SMC and/or HFD interventions in mice 20 weeks of age, corresponding to matured adulthood. The idea behind this was that akin to room temperature housing, younger mice expend a greater proportion of their calories towards growth (48). Neither of these changes was sufficient to cause kidney injury, but the early onset of fibrosis found in these kidneys suggests that they may contribute to accelerating CKD. Another factor we did not examine was using a strain other than the C57BL6/J mice used in the current study. Other mouse strains thought to be more prone to developing CKD, as their genetic makeup influences their susceptibility to illnesses (49, 50). C57BL/6J mice have been demonstrated to be more resistant to developing kidney injury compared to other mouse strains, such as BALB/c or 129/Sv mice (50–52). Nevertheless, C57BL/6J mice are prone to diabetes, insulin resistance, and obesity when fed a high-fat diet, which are significant risk factors for CKD in humans (42, 49, 53). We also focused the current study on the C57BL6/J strain because many transgenic and knockout mouse lines are available on this background.

Obesity-induced CKD leads to malfunction in the mitochondria, thus altering the metabolic program of proximal tubule cells (54). Impaired mitochondrial function has been implicated in the pathogenesis of CKD (55). In addition to impaired ATP generation, reduced fatty-acid oxidation would be expected to promote steatosis, and inefficiency in the electron transport chain would be predicted to increase oxidative stress. Nonetheless, our SMC and/or HFD interventions did not induce these changes. There were also no changes in mitochondrial density or the concentration of respiratory enzymes in isolated kidney mitochondria. HFD feeding appears to increase mitochondrial respiration, consistent with an early adaptive response previously reported (43, 56).

In conclusion, these results highlight the technical difficulty of simulating human chronic kidney disease in mice, especially considering the length of the intervention we utilized in this study. While our insults were not robust enough to reduce GFR, some regions of the kidney showed early signs of fibrosis. Thus, it remains possible that combining SMC and HFD interventions using different strains of mice or with knockout/transgenesis may be useful in modeling human CKD. Mice continue to be widely used in modeling human disease because their genes are easy to manipulate. While we were not successful in developing our CKD model, our findings support the notion changes in diet and physical inactivity may be insufficient to induce CKD. These findings further demonstrate the importance of developing a translatable CKD model to accelerate discoveries of the underlying mechanism of and treatment for human CKD.

## AUTHOR CONTRIBUTIONS

P.C.O. conceived and designed research, performed experiments, analyzed data, interpreted results of experiments, prepared figures, drafted the original manuscript, edited and revised the manuscript; S.T.D. performed experiments and edited and revised the manuscript; D.S. performed experiments and analyzed data; A.D.P. analyzed data; V.L.P. performed experiment; P.S. designed research; M.J.D. and A.S designed research and revised manuscript; N.R. designed research, interpreted results of experiments; K.F. conceived and designed research, interpreted the results of experiments, edited and revised the manuscript, and approved the final version of manuscript.

## ACKNOWLEDGMENTS

This research is supported by National Institutes of Health (NIH) grants DK107397, GM144613, DK127979, and AG074535 to Katsuhiko Funai, DK133271 and HL155345 to Nirupama Ramkumar, DK130555 to Alek Peterlin, Institutional National Research Service Award (5T32DK091317), the National Kidney Foundation of Utah and Idaho to Stephen T. Decker, and American Heart Association grant (AHA 915674) to Piyarat Siripoksup.

---

**Supplemental Table 1:**
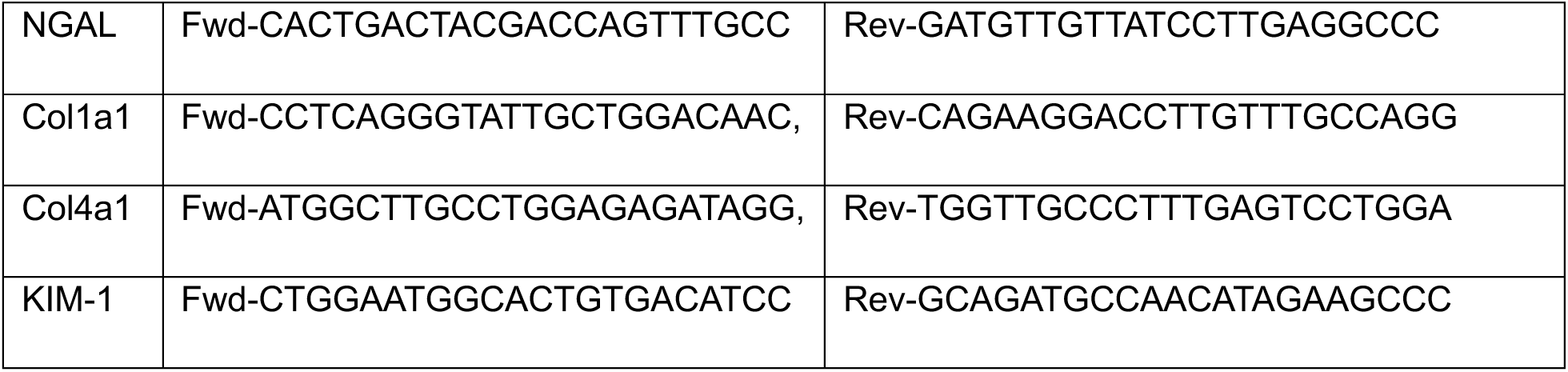
List of primers and their sequences.

**Supplemental Figure 1.**
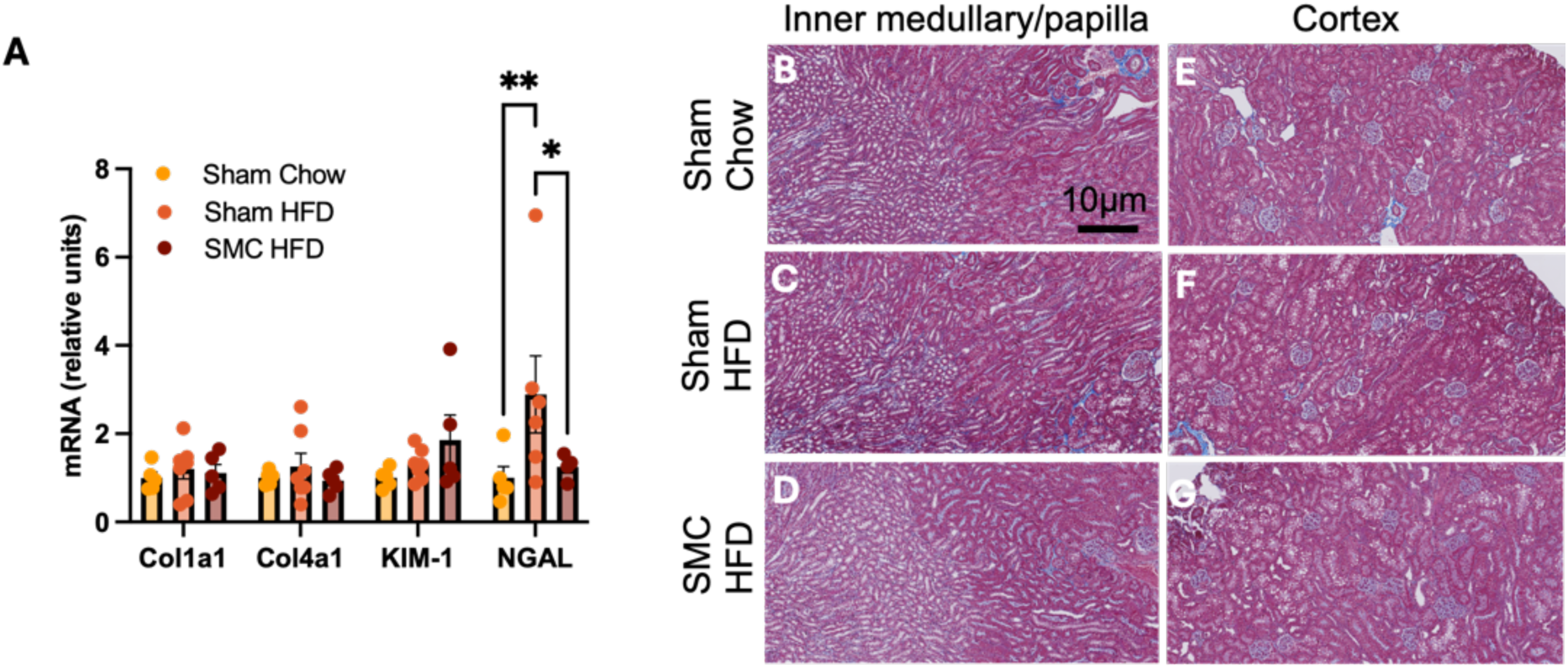
Combined SMC and HFD intervention induces mild kidney fibrosis. (A) Kidney damage markers: collagen I & IV, kidney injury marker-1 (KIM1), and neutrophil gelatinase-associated lipocalin (NGAL) relative expression. Trichrome-stained kidney inner medullary/papilla (B-D) and cortex (E-G) sections from sham and SMC mice fed chow or HFD housed at room temperature. P value =0.05 (P value *<0.05, ** <0.005, *** <0.0005). Images are representative of n=5-10 per group. Magnification at 20x zoomed using Zeiss AxioScan.Z1 slide scanner. HFD ⎯ High-fat Diet; SMC ⎯ Small mouse cage; RT ⎯ Room temperature.

**Supplemental Figure 2.**
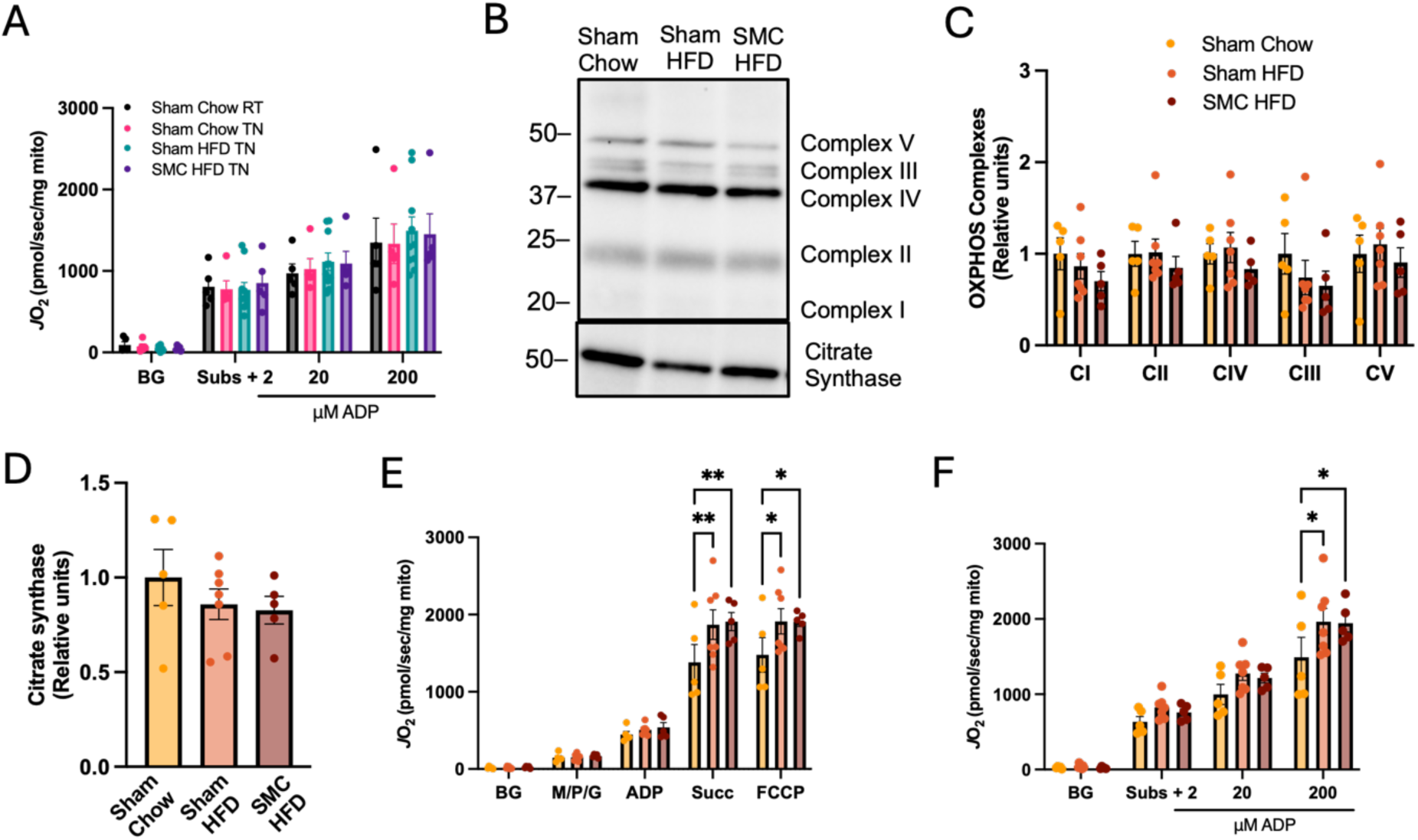
Mitochondrial bioenergetic phenotyping in kidney. (A) O_2_ utilization with ADP titrations (TN), (B) Western blots, and (C) quantifications for OXPHOS complexes I - V and (D) citrate synthase in whole kidney lysate. (E) O_2_ utilization using enriched kidney mitochondria fraction with glycolytic substrates (2 mM ADP, 0.5 mM malate, 5 mM pyruvate, 10 mM succinate, 1 μM carbonyl cyanide-*p*-trifluoromethoxyphenylhydrazone (FCCP) and (F) O_2_ utilization with ADP titrations (RT). n=5-10 per group. HFD ⎯ High-fat Diet; SMC ⎯ Small mouse cage; RT ⎯ Room temperature; TN ⎯ Thermoneutrality.

**Supplemental Figure 3.**
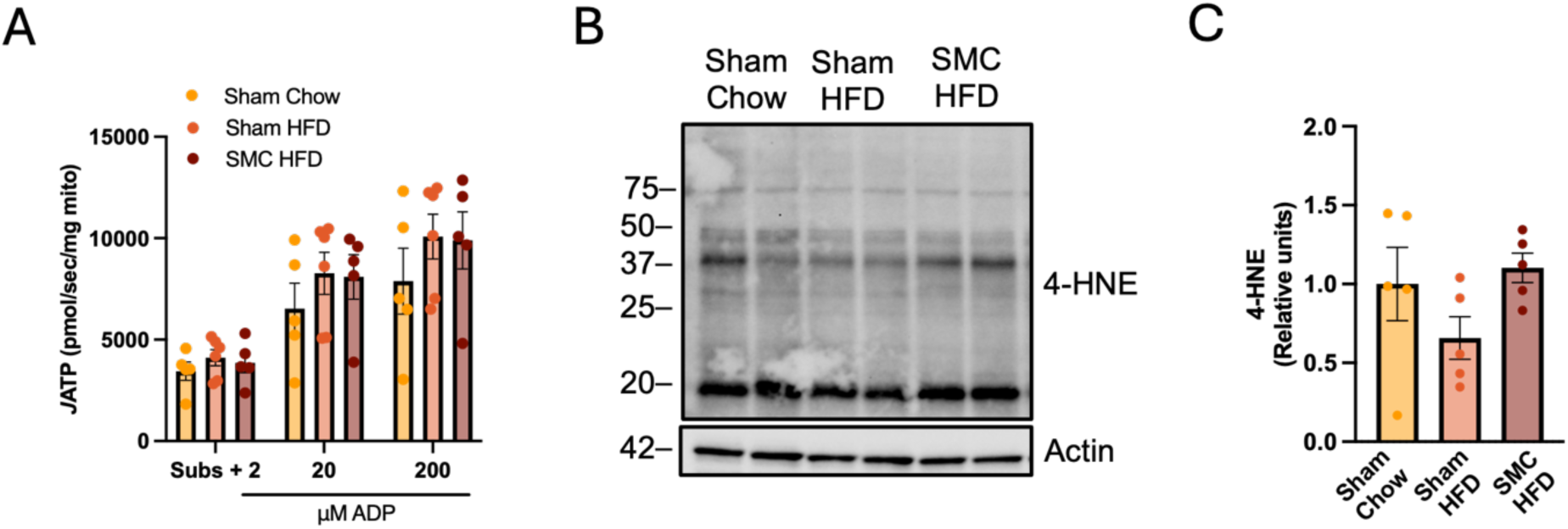
Mitochondrial ATP production and electron leak. (A) ATP production with ADP titrations using enriched kidney mitochondria fraction. (B) Western blots and (C) quantification for 4-hydroxynonenal (4-HNE) and actin in whole kidney lysate of sham and SMC mice at room temperature, n=5-10 per group. HFD ⎯ High-fat Diet; SMC ⎯ Small mouse cage; RT ⎯ Room temperature; ADP-Adenosine 5’-diphosphate.

